# Pupil responses to social stimuli are associated with adaptive behaviors across the first 24 months of life

**DOI:** 10.1101/2025.05.30.656675

**Authors:** Rebecca Grzadzinski, Raymond S. Carpenter, Josh Rutsohn, Alapika Jatkar, Kattia Mata, Ambika Bhatt, Maria M. Ortiz-Juza, Madison R. Dennehey, Donna Gilleskie, Jed Elison, Nicolas Pégard, Jose Rodriguez-Romaguera

**Author notes:** Co-first Authors (Rebecca Grzadzinski, Raymond S. Carpenter). Address correspondence to: Rebecca Grzadzinski, PhD, Assistant Professor, University of North Carolina, Chapel Hill, NC, Jose Rodriguez-Romaguera, PhD, Assistant Professor, University of North Carolina, Chapel Hill, NC.

## Abstract

**Background:** Pupil changes in response to well-controlled stimuli can be used to understand processes that regulate attention, learning, and arousal.

This study investigates whether pupil dynamics to social stimuli are associated with concurrent adaptive behavior in typically developing infants. To accomplish this, we developed and assessed pupillary responses to Stimuli for Early Social Arousal and Motivation in Infants (SESAMI).

**Methods:** A sample of forty-six typically developing children aged six to twenty-four months were exposed to SESAMI. Infants were presented with either dynamic social faces or non-social stimuli that controlled for luminance, motion, and auditory exposure. A multi-level mixed effects model was used to fit pupillary response functions (PRFs) that measure the change in pupil size over time as the infants fixate on either a socially dynamic face or a non-social control. This model produces separate social and non-social PRFs for both the population and each individual. An average individual deviation score from the population PRF was calculated separately for social and non-social trials yielding the pupil response index (PRI). Vineland Adaptive Behavior Scales (VABS) were regressed on social and non-social individual PRIs while controlling for age and average fixation time. We tested whether the social PRI was a statistically significant predictor of adaptive behavior by comparing the model predicted VABS scores with observed scores.

**Results:** An increase in PRI to social stimuli was significantly associated with better adaptive behaviors in typically developing children between 6 and 24 months of age.

**Conclusions:** SESAMI combined with pupillometry and multi-level mixed effects modeling, provides a novel and scalable framework for quantifying individual differences in pupil changes in response to social stimuli relative to a population-level baseline. By demonstrating that pupil response indices during social fixations predict adaptive behaviors, we lay the foundation to test how these measures may help identify infants with intellectual and developmental disorders.

## Background

Changes in pupil size are widely used in preclinical science as a metric that represents neurobiological states, and behaviors associated with underlying neural circuits^1–5^. Pupil size (dilation and constriction) is guided by underlying sympathetic and parasympathetic mechanisms and changes reflexively and dynamically in response to luminance shifts as well as in response to cognitive load or attentional activities^5^. Though the exact meaning of pupil changes is not fully understood, preclinical work often refers to these dynamic changes as “arousal” processes^2,3^. Both non-human and human literature has correlated pupil size with other physiological arousal metrics including heart rate and skin conductance^6–8^. Pupil changes have also been linked to neural activity in the cortex, thalamus, superior colliculus, and the locus coeruleus^3,9–11^. As such, pupillary dynamics has been suggested as an indirect strategy to assess underlying neural processes associated with human behaviors such as cognition, sensory reactivity, and learning and memory processes^12,13^. However, there is a scarce amount of literature exploring pupil changes to specific stimuli in humans beyond reflexive responses to luminance changes (pupillary light reflex)^5^. This is likely due to the variety of internal and external processes that can impact pupil dynamics (such as environmental luminance, attention, and cognitive load) which are often opposing and/or not fully understood. Therefore, assessments that assess pupil dynamics in response to well-controlled stimuli (of static luminance) allow us to extract pupil dynamics that are more related to attention and cognitive load processes, which in turn provide key insights into a variety of typical and atypical processes. This insight has inherent application to intellectual and developmental disorders such as autism spectrum disorder (ASD), Down syndrome (DS), and attention deficit hyperactivity disorder (ADHD), as well as neural degenerative processes such as dementia and parkinsonism^14,15^.

In humans, research has linked pupil changes in response to stimuli with reward and social learning processes. For example, infant pupils dilate while viewing, and in anticipation of, a video of their mother compared to a video of a stranger, highlighting the pupil as a metric to quantify the value of a social stimuli for infants^16^. Increased pupil size has also been linked to the anticipation of earning monetary rewards in adults, suggesting an association with reward processing and learning^17^. Beyond pupil dynamics, infant gaze following has been linked to later vocabulary growth and language development^18,19^. Gaze following has also been linked to other physiological metrics such as elevated heart rate during eye contact^20^. Decreased heart rate variability and respiratory sinus arrhythmia (RSA) during play has also been linked with poorer social communication skills later in life^21^. Further research is required to disentangle the relationship between pupil dynamics and other physiological arousal metrics and later social skills. In the neurodevelopmental literature, some evidence indicates that autistic children, or those at elevated familial likelihood for ASD, show differences in their pupil dilation in response to changes in dark-light stimuli (PLR) compared to controls^22,23^. For example, a stronger PLR in high likelihood children is linked with more severe ASD symptoms and worse functional outcomes, such as cognitive delays and attention difficulties^24^. Beyond PLR, children with ASD have been found to have no change in pupil size while watching an actor’s emotional reaction to opening a box^25^. Another study found that 9-month-old infants at elevated likelihood of ASD had a larger average pupil size while looking at emotional faces compared to infants at low familial likelihood for ASD (no sibling with ASD); this increased pupil response at 9 months was associated with worse social communication skills at 18 months^26^. Unfortunately, the diagnostic outcomes were not presented^26^, and average pupil size may represent a diluted or inaccurate representation of a more complex, temporally dynamic process. Outside of ASD, increased pupil dilation in response to an oddball paradigm was seen in DS, suggestive of increased cognitive effort^27^. Children with ADHD had a decreased pupil size while completing a visuo-spatial working memory task compared to typically developing children; this difference in pupil size was no longer present in ADHD when medication was taken^28^. Overall, studies that robustly evaluate pupil dynamics in response to well-controlled social stimuli, across typical and atypical samples, and that link these findings to diagnostic outcomes and functional behaviors are necessary to elucidate the utility of this metric.

The potential benefits of pupillometry as an objective assessment tool cannot be understated. Changes in pupil size in response to stimuli (pupil dynamics) are easily obtained, noninvasively across development and age. Specific skills or a written/spoken response are not required. With clear links to physiology, neurobiology, and application to preclinical studies, the pupil is an ideal candidate to target in studies of those with diminished or limited capacity, such as infants or those with neurocognitive deficits. In this study, we evaluate the relationship between pupil dynamics in response to well-controlled social stimuli and adaptive functioning in typically developing children. We hypothesize that unique pupil dynamics during fixation to social stimuli will be associated with better adaptive skills in typically developing children.

## Methods

### Participants

Forty-six typically developing infants, aged 6 to 24 months old, were enrolled. Fifty-eight percent of participants were female, and 75% of those who provided race information were white (race information was missing for 20% of participants). All study procedures were approved by the institutional review board of the University of North Carolina at Chapel Hill, and all parents/guardians provided written informed consent to participate. Participants were recruited from the community, such as pediatrician offices, public flyers, and parenting groups on social media. Inclusionary criteria included being between 6 and 24 months of age, being at low likelihood for developing autism (i.e., no first-degree relative with autism), and living in a household where English is the primary spoken language. Exclusionary criteria included having a known genetic condition, significant motor delays, or known visual or auditory impairments.

### Study procedures

The in-person assessment included an eye-tracking battery and a parent interview using the Vineland Adaptive Behavior Scales, Third Edition (Vineland-3^29^). Eye-tracking was completed upon arrival at the testing site to maximize attention, unless the child required time to settle before attempting eye-tracking. Parents completed surveys including a brief medical history.

### Vineland-3

The Vineland-3^29^ is a parent interview administered by a trained assessor to evaluate adaptive functioning in individuals across the lifespan. The VABS provides an Adaptive Behavior Composite (ABC) standard score (mean=100, SD=15), that is an overall impression of the child’s adaptive skills. The Vineland-3 also provides standard scores for areas of Communication, Socialization, Daily Living, and Motor Skills, though these are not included in analyses due to the limited scope of this work and sample size.

Within the Communication domain, parents answer questions regarding the child’s understanding of words and messages, and early expressive behaviors such as vocalizations and gestures the child expresses independently. The Daily Living domain assesses the child’s eating and drinking habits, and their independence in routines such as hygiene and dressing. The Socialization domain gathers information on early social behaviors including smile responses, engaging in play with others, and showing interest in peers or familiar people. The Motor skills domain evaluates foundational physical ability, such as sitting or standing, as well as fine-motor coordination, and object handling skills. For this study, we use only the VABS ABC scores due to the sample size. Future analyses will explore domain-level relationships.

### Eye Tracking Procedures

The eye-tracking battery lasted approximately 15 minutes and consisted of five total stimuli (two calibration tasks). The battery began with a five-point calibration and validation model, using corner and mid-screen focal points respectively. The SESAMI paradigm was preceded by two other stimuli videos (not part of this manuscript), depicting both real and avatar human forms. The session concluded with a final five-point validation calibration.

Infants were either securely seated in a highchair (50%) or their parent’s lap (50%). In the cases of infants sitting in their parent’s lap, parents wore infrared (IR) blocking glasses to ensure only the infant’s eyes were captured. The paradigm was conducted in a well-lit room (260 lux) with parents and the assessor in close proximity, but with instruction not to directly engage with the child during the paradigm. Measurements of gaze duration, pupil dilation, and fixation were all obtained via the Tobii Pro Lab Full Edition, version 8.0. Each stimulus was presented on a 24-inch screen with LCD binocular eye tracking, capturing at a rate of 600-Hz and a resolution of 1920 x 1080. Infants were positioned approximately 60-cm away from the center of the Tobii Pro Spectrum monitor. From the infants’ perspective, the display covered about 47° horizontally and 28° vertically of visual angle on average. Infants were unable to view the assessor’s Tobii Pro Lab screen tracking eye movements.

### Stimuli for Early Social Arousal and Motivation in Infants (SESAMI)

SESAMI was presented as a series of video clips with a different female model shown in each block. The average block (static non-social, dynamic non-social, static social, dynamic social) was 23 seconds in length. Each block has a static and dynamic component as well as a social- and non-social component, shown in the order: static non-social, dynamic non-social, static social, and dynamic social. The dynamic social component consisted of a female model saying a warm greeting (“You’re so precious”, “It’s so good to see you!”). The non-social component was created by inverting, pixelating, and distorting the social model video to remove any identifiable biological motion while maintaining luminance. The audio file was replaced by pitch-matched piano tones generated with the ScoreCloud software, to match any possible arousal caused by auditory stimulation. To obtain static counterparts, the first frame of each dynamic portion was held for 5 seconds. In total, there were nine blocks, with the order of blocks shown randomized into groups A and B, which were alternated across participants. The total paradigm was three minutes and twenty-seven seconds.

### Data Preparation

Pupil size measurement was extracted from the right eye if valid or the left eye if the right eye was invalid to produce one pupil size measurement. Ninety-four percent of the measurements came from the right eyes, and six percent came from the left eyes. To remove blinks and artifacts, we applied a variance-based thresholding method^30–33^. We first normalized the data using Min-Max normalization to standardize measurements across participants^34^. After normalization, a square root transformation was applied to address the left skew introduced through normalization^35,36^. Rolling variance was computed over a time-based window to capture short-term fluctuations. We also computed the rate of change between consecutive time points (in min–max % per ms) and flagged any change greater than 2.5 min–max %/ms. Both metrics were necessary because rolling variance alone could miss rapid but subtle drops, and rate of change alone might fail to account for larger artifacts with slower transitions. The rolling variance threshold was set at the 75th percentile of all variances, reflecting the upper range of expected physiological noise. Anything that fell outside of the range for the rolling variance or rate of change filters was flagged and a 50-millisecond buffer was applied around flagged centers to remove artifacts^30^. The Min-Max of pupil size was only used during data preparation and visualizations in Figure 1, the pupil size in millimeters was used for all subsequent statistical analysis. After the artifact and blink removal forty-five participants remained for further analysis.

**Figure 1.**
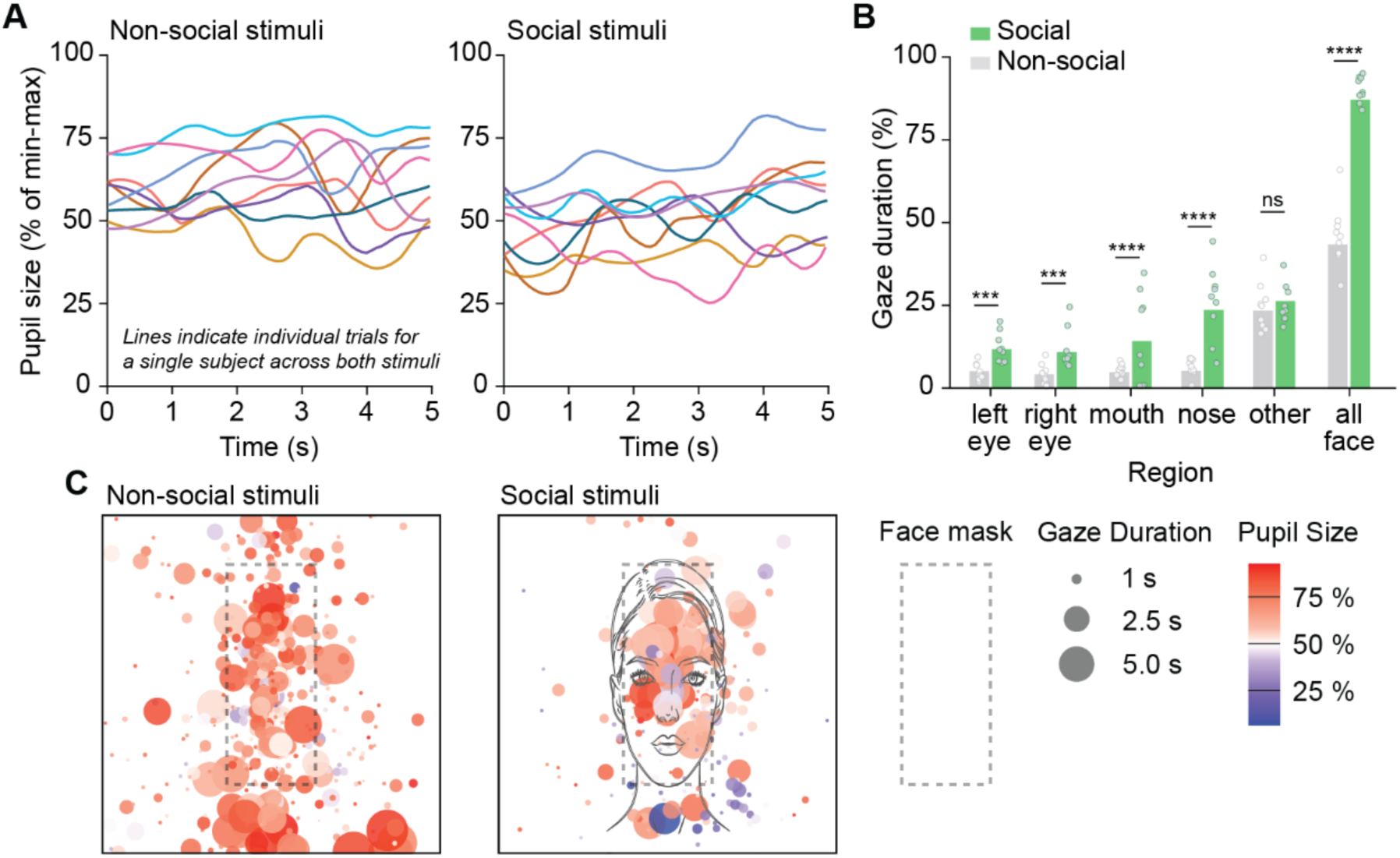
Infants fixate more on social stimuli than non-social stimuli and exhibit diverse pupillary responses to social stimuli. **(A)** Representative raw min-max normalized pupil size dynamics from a single infant across multiple trials with non-social (left) and social (right) stimuli. Each colored line represents an individual trial, showing fluctuations in pupil size (expressed as a percentage of the subject’s total min-max range) over time. **(B)** Comparisons of the percentage of gaze duration directed at different facial regions between social and non-social stimuli. Each dot represents the average gaze duration across participants for a specific trial. However, the statistical analysis and error bars were computed based on the participant level averages (n=45). Statistical significance is indicated as follows: ***p<0.001, ****p<0.0001, ns = not significant. **(C)** Heatmaps overlaying gaze duration (circle size) and pupil size (color intensity) on face stimuli. Warmer colors (red) indicate larger pupil sizes, while cooler colors (blue) indicate smaller pupil sizes. Larger circles represent longer gaze durations. Data are shown separately for non-social (left) and social (right) stimuli for all participants across one trial.

We down sampled the data to 40 Hz by averaging pupil size measurements, doubling the recommended minimum sampling rate of 20 Hz to capture physiologically natural pupillary dynamics^33^. We calculated the point luminance from the image from an area around where the eye was fixated with ten percent of the entire screen size^37^. The point luminance captures the average luminance coming from the screen and emanating to the eye in a single scalar. The ambient luminance, coming from the surrounding environment, is controlled in the mixed model through random effects. We did not analyze differences between dynamic and non-dynamic portions of the trial, as this was beyond the study’s scope. To control for differences between dynamic and static scenes, we added a variable that flags each scene as dynamic or static to the pupil-response model.

### Pupillary Response Functions

We extracted a social and non-social pupil response function from the data at both the individual and population levels. We define the pupil response function as the fitted curve of pupil size measured from the start to the end of a continuous fixation. This curve captures how the pupil changes over time in response to a stimulus. Figure 1A depicts pupillary dynamics across the 9 trials. In our study, the response onset is marked by the moment the subject fixates on a social face region. The Tobii I-VT Fixation Filter was used, which classifies fixations and saccades based on the velocity of eye gaze shifts^38^.

To define social face regions, we mapped areas surrounding the face, each eye, the nose, and the mouth for both the social trials and their corresponding areas in the nonsocial trials. These regions spanned about 5.2°×2.6° of visual angle for both eyes, 3.8°×3.5° for the nose, 6.4°×3.0° for the mouth, and 13.0°×18.2° for the whole face. Figure 2C illustrates the borders of the social face regions (face mask) used for both the control and stimulus trials. Figure 1B shows gaze duration (fixation) in face regions across social and nonsocial trials.

**Figure 2.**
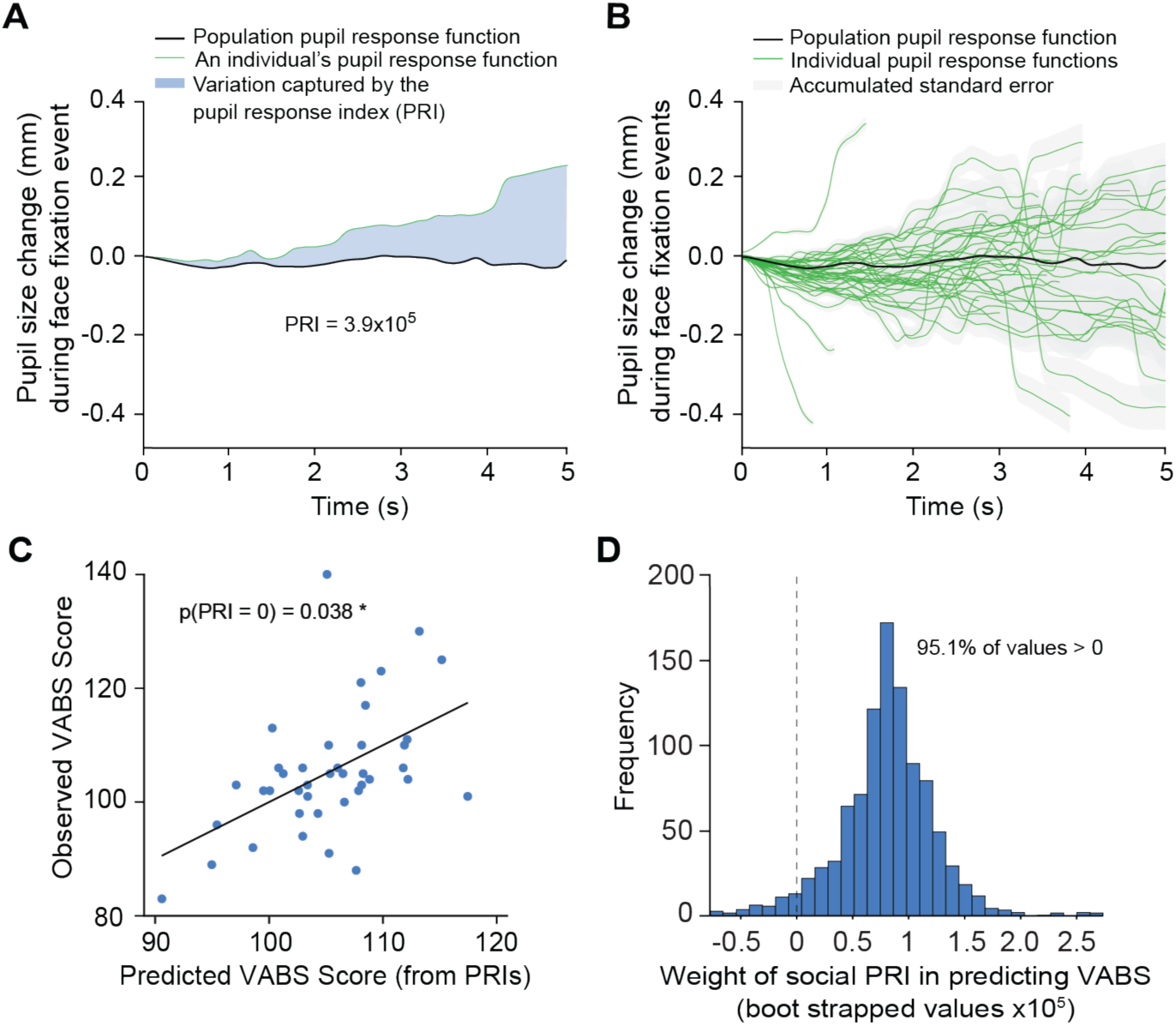
Estimated pupillary response deviations allow for the prediction of adaptive behavior scores in infants. **(A)** The shaded area captures the deviation between the average pupil response function and an average pupil response index for one individual over the first 5 seconds. **(B)** Population-level and individual estimated pupil response functions during the social stimulus trials generated by fitting a multi-level mixed effects model. Shaded area represents accumulated standard error, the green lines show individual pupil response functions, and the black line shows the population pupil response function. **(C)** A scatterplot depicting the relationship between the predicted and observed VABS scores. Each point represents an individual, with predicted scores on the x-axis and observed scores on the y-axis. The black line represents a regression of observed scores on the predicted values, serving as a visual check on how well predicted values align with observed values. The reported p-value, “p(social PRI = 0) = 0.034”, tests whether the social PRI significantly contributed to the model’s predictive ability. This indicates that PRI captures meaningful individual variation in social pupillary responses that predict observed VABS scores. **(D)** Distribution of 1000 bootstrapped social PRI weights for predicting VABS scores, generated by resampling participants with replacement. The dashed line represents the null hypothesis (social PRI = 0). Since 95.1% of bootstrapped values are above 0, we reject the null hypothesis. This further supports the conclusion that social PRI meaningfully predicts VABS scores.

### Data Analysis

Prior to modeling, we examined whether infants allocated more time to social stimuli compared to non-social stimuli. Previous studies have shown that typically developing infants prefer a social stimulus over a non-social stimulus^39,40^. To validate this within the SESAMI paradigm, we computed the average fixation time in social blocks and contrasted it with the total fixation time in non-social blocks. We performed this test using a repeated-measures t-test to account for within-subject differences in fixation behavior. Comparisons were made separately for each corresponding facial region in the social and non-social blocks. These regions included the mouth, nose, left eye, right eye, and “other” regions surrounding these features, and an “all-face” region encompassing them all. The statistical analyses and error bars were calculated on participant-level averages. Forty-five participants were used for this analysis.

Only pupil measurements recorded during fixations on the designated facial regions (non-social control or social stimulus) were included for the next portion of analysis. Any data recorded outside those fixations were excluded as they did not reflect the relevant pupillary events. Only the face designation was used here. The mouth, nose, and eyes were all considered part of the face. Twenty-four percent of observations were not considered face fixations and were dropped. In total, 5 more subjects and 7.5 percent of all trials were dropped due to not having any face fixations for both the non-social and social trials.

After removing data points during periods of blinks, artifacts, and no face fixation, 139,338 data points (32 % of the initial 427,802) remained for the mixed-effects analysis. Observations came from 40 participants ages 6 to 22 months, encompassing 570 trials, and 3,453 distinct face fixations. Participants showed a mean fixation duration of 1,666 ms (SD = 2,596 ms), made an average of 93.1 fixations across all trials, and contributed an average of 7.6 social trials and 7.4 non-social trials.

A two-stage model was used to characterize pupil response functions and link them to adaptive behavior outcomes. A multi-level mixed effects model was used in the first stage to fit all the pupil response functions. Mixed models have been shown to successfully detangle spatial, object, and atypical arousal responses linked to ASD in adults^30,41–43^. Baseline pupil size and ambient luminance was captured by including a trial-level random intercept, so any constant lighting or average pupil size differences across trials are absorbed into each trial’s intercept rather than contaminating the pupil-response estimates. Including the random effects ensures that time-invariant participant characteristics and environmental differences are properly accounted for, which generates a robust population average pupil response function.

Including trial-level random slopes for the pupil response in the first order allows for an individual pupil response index (PRI; captured through the Best Linear Unbiased Predictions (BLUPs) of the random effects). This random slope, or PRI, captures the individual variability in the dynamic pupil response for each trial. The average social and non-social PRI relating to the pupil response function was derived for each participant, creating two scalars for everyone that captured the deviation from the average pupil response function for both the social and non-social PRIs. The individual average and population social pupil response functions can be seen in figure 2A.

The fit of the pupil response function is measured by the adjusted R^2^ and Root Mean Squared Error (RMSE) of the first stage multi-level mixed effects model. Forty participants were used for modeling, as four were excluded due to an insufficient number of face fixations across trials. By incorporating fixed and random effects, our model accounts for population-level factors, such as the average pupillary response, and individual- and trial-level variability, using random intercepts to model the baseline pupil size and random slopes to capture trial-specific fluctuations in pupillary response and luminance response^44^.

To determine whether these individualized pupil deviations carried predictive value for adaptive behavior, we regressed VABS scores on each participant’s social and non-social PRIs, along with age and fixation time as controls. We evaluated whether the Social PRI coefficient differed significantly from zero by computing its t-statistic and p-value in STATA^45,46^. In addition, a percentile-based bootstrap with 1,000 resamples was performed to generate a distribution of the Social PRI coefficient and test whether 95% of these coefficients were above 0.

## Results

### Infants fixate more on social stimuli than non-social stimuli during SESAMI

To validate SESAMI we measured pupil dynamics in typically developing infants and first examined whether our cohort exhibited a preference for social stimuli compared to non-social stimuli^39,40^. We quantified gaze behavior by measuring the percentage of time infants fixated on specific facial regions across all social and non-social stimulus trials. For each trial, gaze duration was averaged across all infants. Our results showed that, on average, infants showed significantly longer fixation to the social condition for the left eye (p < 0.001), right eye (p < 0.001), mouth (p < 0.0001), nose (p < 0.0001), and the overall face (p < 0.0001), whereas the “other” facial region was not significantly different between blocks (Figure 1B).

### Pupillary Responses to Social Stimuli Captured Through Mixed Effects Modeling

We used a multi-level mixed effects model in the first stage of our two-stage model to establish a robust framework for predicting pupil dynamics over time, generate both individual and average pupillary response functions, and capture deviations of individual responses from the average (PRIs; Figure 2A). The multi-level mixed effects model produced an adjusted R^2^ of 0.92, meaning the model that fits the pupillary response functions accounts for 92% of all variation observed in pupil size. It also produced a RMSE of 0.15, meaning each prediction was on average 0.15 millimeters off from the true pupil size.

### Individual Social Pupillary Response Functions Predict VABS scores in infants

To evaluate whether the individual deviation from the average pupillary response (might serve as a marker for predicting adaptive behavior scores, we regressed VABS scores on the individual average social PRIs, individual average non-social PRIs, average fixation time, and age. Figure 2A illustrates the shaded area between an individual’s social pupil response function and the population average, which defines that participant’s social PRI that we used in the second stage regression. The social PRIs were statistically significant with a p-value of 0.034, indicating a relationship between pupil dynamics during social fixation and VABS (Figure 2C). The bootstrap test was also statistically significant to the 95% confidence level (p<0.05). 95.1% of all coefficients were above 0 (Figure 2D). This second-stage regression produced an adjusted R^2^ or 0.19 and an RMSE of 10.1.

## Discussion

This study is the first to evaluate the relationship between pupil dynamics and adaptive behaviors in typically developing children between 6 and 24 months of age. We used novel mixed modelling to evaluate pupil size as a temporally dynamic response to well-controlled social and non-social stimuli. This technique provided a pupil response function (PRF) which was then used to create an individual pupil response index (PRI)-- a value representative of the individual deviation from the mean. We found that larger PRI (more pupil dilation) in response to fixation on the social face was associated with better adaptive behavior scores. This finding replicated when using a bootstrapping method with more than 95% of the 1000 iterations yielding significant results. Overall, these findings highlight the value of pupil dynamics as a relevant biomarker of skills in infants and toddlers. Further studies are required to fully elucidate the potential of pupil dynamics and to better evaluate which underlying constructs differentially impact it. Pupillometry is an ideal target for study in diverse populations as it is easily acquired across developmental levels and requires minimal resources^47^. To date, pupil dynamics have been interpreted as based in arousal, attentional, sensorial, effortful, or learning processes^12,16,17,24–28^. Continued studies using well-controlled stimuli with robustly phenotyped and longitudinal samples can help distinguish these processes.

We also confirm findings from other work that typically developing children look at social stimuli more than nonsocial stimuli (see Figure 1B)^48–50^. Research also indicates that those with or at high likelihood for developing autism may fixate on the face less than those with typical development^51^; while this work only includes those with typical development, future work will need to examine the differences in baseline fixations across groups. In this work, the average time spent fixating on the face was not predictive of VABS scores suggesting that social fixation, which may reflect attention processes, and are distinct from pupil dynamics described in this study which have a unique relationship with adaptive skills. Attention processes, which may or may not be adequately defined in this work (fixation time) require further exploration to understand the relationship with pupil dynamics. For example, future work should explore the number of fixations as well as combine fixation information over time, despite the interruption of saccades. Another novel addition to this work includes the quantification of individual pupillary response to the localized luminance reflecting from the fixation location. By quantifying the local luminance and accounting for the constant environmental luminance through mixed effects, we can remove the pupillary response that is specific to luminance from the pupillary response that is specific to social fixation.

Our findings have clinical implications for neurodevelopmental disorders (NDDs), such as ASD and Down syndrome (DS) as well as neurodegenerative disorders such as Parkinson’s and Alzheimer’s disorders. With the use of SESAMI, we were able to predict adaptive behaviors with high accuracy using a model that combines pupil dynamics and chronological age. This metric has high potential as a biomarker for adaptive, and potentially cognitive, skills that can be quickly, easily, and validly acquired across ages, capacity, and development. For example, DS research is in dire need of valid measures of adaptive and cognitive skills that can accurately quantify the heterogenous profiles that are characteristic of the lower bounds of traditional standardized testing^52–54^. PRI might provide clinicians and researchers with a useful tool to estimate cognitive or adaptive skills in those with limited verbal or diminished capacity.

Our findings may also apply to diagnostic outcomes, such as autism or other neuropsychiatric conditions. For example, autism spectrum disorder (ASD) is a pervasive NDD characterized by deficits in social communication and the presence of restricted and repetitive behaviors^55^. Symptoms emerge gradually over the first few years of life and is preceded by a period of seemingly typical development —a period during which symptoms of ASD are either absent or only present in their very mildest forms^47,56^. Very early markers that are predictive of ASD outcomes, particularly at an individual level, are critical to facilitate access to very early intervention windows—when treatment may be more effective^47,57^. PRIs described in this manuscript may provide just such a biomarker of later ASD diagnoses.

As a group, children who go on to develop ASD show behavioral differences from those who do not develop ASD during this presymptomatic period (e.g., lower cognitive ability). However, on an individual level, an infant who goes on to receive a diagnosis of ASD is behaviorally indistinguishable from one who does not^58–60^. While observed behavior has had limited utility in individual diagnostic identification before 24 months of age, neurobiological differences by 6 and 12 months of age have shown more promise, particularly in high familial likelihood (sibling) samples. For example, Hazlett et al.^61^ found that cortical surface area measurements at 6 and 12 months of age were predictive of ASD diagnosis at 24 months of age, on an individual level, with a positive predictive value of 81%, consistent with the accuracy of most established diagnostic tools (e.g., ADOS-2^62^). Similarly, Emerson et al.^63^ found that functional connectivity patterns across the brain of 6-month-olds could accurately predict which infants would go on to develop ASD with a positive predictive value of 100%. The clinical impact of these findings is immense, but the utility is limited by the lack of replication as well as limited access to MRI in community samples. Identification of low-cost, clinically meaningful presymptomatic markers that can be linked with underlying neurobiology continues to be of critical importance. The pupillary response function may be just such a marker that can be linked to underlying neurobiology and ascertained in a low-cost manner.

This work should be considered in light of several limitations. The children in this study were assessed at a single time point. Therefore, our results cannot be interpreted as developmental, though we hope to examine this in future study samples that are seen longitudinally. It is likely that age and developmental processes are likely to affect pupil dynamics^64^. As such, longitudinal samples should include multiple timepoints within 6 and 24 months to thoroughly disentangle the relationship between these pupillary dynamics and changes in age/development. We used a parent report metric of adaptive behavior -- the VABS. While VABS is a commonly used and well-validated tool for measuring adaptive skills, like any parent report measure, it may have a bias as it is not an objective indicator of skills. All the children in our sample were typically developing at the time of data collection and predominantly White. Therefore, these findings require extension to new, more diverse samples to truly elucidate the utility of this paradigm. Lastly, extracting useful, interpretable, and dynamic pupil information from time-series data is complex. This process requires making statistical assumptions and setting data inclusion cutoffs that may not accurately reflect larger populations. We believe our methodology makes the best possible assumptions considering the available data, though replication of our findings will be critical for further validating our results.

## Conclusions

This study highlights the potential of pupil dynamics as an accessible, objective measure with direct links to that predicts adaptive functioning in children between 6 to 24 months of age. By isolating individual differences in pupillary responses through mixed modeling, we demonstrated that larger pupil dilation in response to social stimuli correlates with better adaptive behavior scores. Our findings lay the groundwork for future research to evaluate whether early pupillary response patterns might serve as a clinically useful metric of adaptive skills that can be used across neurodevelopmental and neurodegenerative disorders.

## List of abbreviations

ASD: Autism spectrum disorder
BLUPs: Best Linear Unbiased Predictions
PRF: Pupil response function
PRI: Pupil Response Index
RMSE: Root Mean Squared Error
SESAMI: Stimuli for Early Social Arousal and Motivation in Infants
VABS: Vineland Adaptive Behavior Scales

## Declarations

### Ethical Approval

Study procedures were approved by the Institutional Review Board at the University of North Carolina at Chapel Hill (IRB #21-1704).

### Consent for Publication

All authors have approved the manuscript for publication.

### Availability of data and materials

Available upon request to the corresponding authors.

## Author Contributions

Authors RG, RC, JRR, NP were involved in project conceptualization and development, task development, data analysis, and manuscript preparation. RG, KM, AJ, and AB were involved in data collection, organization, and cleaning. AJ was involved in participant recruitment and AJ, KM, and AB were involved in project management. MRD assisted with major revisions to the manuscript. JR, MOJ and KD were involved in data analysis. JE developed original task stimuli.

## Acknowledgments

Special thanks to Drs. Heather Hazlett, Joseph Piven, and Georgina Lynch for their support of this project. The authors are grateful for the contributions of Dr. Kevin Donovan, who contributed to initial conceptualizations of this project.

## Supplemental

### Decision Not to Use Interpolation

Interpolation was avoided to prevent the introduction of artificial data points that could distort physiological patterns and because it was unnecessary based on the chosen model. Instead, we employed a set of polynomials to model the temporal dependencies in the data without the need to include lag variables or conduct interpolation. Furthermore, the mixed-effects model inherently accommodates individual variability and unbalanced data, eliminating the need for interpolation^65,66^. Linear interpolation was also tested with the two-stage model, which did not meaningfully alter the results.

### Model Design

The multi-level mixed effects model is specified as follows:

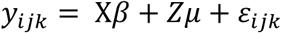

where *y_ijk_* represents the observed pupil size for participant i, event j, and observation k. Х*β* represents the fixed-effects component. This term captures systematic patterns in pupil size by fitting an average pupillary response function. The average pupillary response function is modeled as a polynomial function of time, where time is measured in milliseconds from the start of a fixation. The degree of the polynomial and its interactions were optimized to maximize the adjusted R² for predicting VABS scores in the final model. A third-order polynomial was used to fit the temporal component of the pupillary response function in the multi-level mixed effects model. The binary variable representing the shift between the static and dynamic portions was used as a control variable. Interaction between the binary dynamic indicator variable and each order of the pupillary response function were also added as controls. Additionally, fixed effects control for all time-invariant participant characteristics by capturing the systematic, population-level, relationship between the predictors and pupil size. Thus, holding constant any stable, unchanging participant attributes that could confound these effects^67^. This is achieved by subtracting each participant’s average from each observation, thereby isolating within-participant variation. In other words, the fixed portion estimates how the pupil varies with time on average, independent of participant-specific traits that do not fluctuate over the course of time.

The random-effects portion of the model, Zμ, accounts for individual and event-level heterogeneity by incorporating random intercepts and slopes for each trial nested within participants. Each observation includes random intercepts for each trial (nested within each participant) and random slopes for luminance and fixation time for each trial. These slopes and intercepts are combined with the fixed effects, Х*β*. This may help to control for environmental and contextual factors like luminance, mood, or presentation differences between each trial^44,67^.

We used the Restricted Maximum Likelihood (REML) approach to optimize our model by minimizing the sum of squared deviations at each level and to obtain estimates of the variance components associated with both random intercepts and slopes^68^. The random intercepts and slopes are derived from the model-estimated variance and the residuals of individual trials to estimate each trial’s deviation from the overall average. This procedure ensures that the extra variability introduced through each unique slope doesn’t distort the overall fixed-effects estimates, and it helps prevent overfitting due to the small sample size^69^.

After the model was built, we used Best Linear Unbiased Predictions (BLUPs), referred to as the pupil response index (PRI) in this study, to estimate the deviations of individual observations from the overall model predictions. The PRI leverages the estimated structure of the random effects and the observed data to calculate the most likely deviations for each random intercept and slope. These predictions account for both individual-level and event-level variability while remaining unbiased under the model’s assumptions^45,67^. By capturing these deviations, the PRI allow us to quantify how each observation differs from the population-level estimates provided by the fixed effects.

Each participant is assigned a PRI measurement for each trial corresponding to the fixation time variable. This captures the deviation of their pupillary response function from the trial average (See Figure 3A). For each participant, half of the PRIs corresponds to social trials, and the other half correspond to non-social trials. The average PRI measurement per participant was then taken for both the control and stimulus trials.

To measure whether the average PRI for stimulus trials captures social pupillary information, a VABS composite score was regressed on both the average control PRI and the average stimulus PRI per participant in the second-level regression. The composite score comprises assessments of communication, social interactions, motor skills, and daily living skills. Below represents the model structure used.

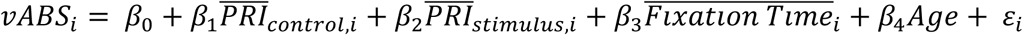

Where i is the index per participant, *ε_i_* represents the error term, *β*_0_ is the model intercept, both *β*_1_ and *β*_1_ are the slopes for both the average control and stimulus PRIs respectively. Additionally, *β*_3_ and *β*_4_ capture the effects of the control variables: average fixation time and age.

To determine whether the average stimulus 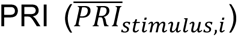 explains a meaningful variation in developmental outcomes, we evaluated the significance of *B*_2_ within the second-stage regression. A t-test was used to test our hypothesis at the 95% confidence level^45^. In the Second-Level regression, various controls were examined to strengthen our tests of *B*_2_. Age was the only control found to be statistically significant in predicting VABS. We retained the average face fixation time as a control variable, even though it was not statistically significant, to bolster our claim that observed association between the stimulus PRIs and developmental outcomes is specific to social pupillary responses rather than merely reflecting overall attention or fixation behavior.

We used bootstrapping to obtain a more robust measure of whether our PRI captures variance related to social behavior scores. We repeated the process of constructing the multi-level mixed effects model, extracting the PRIs, and using them in the second-level regression to obtain 1,000 separate estimates of *B*_2_. Participants were randomly sampled with replacement. This approach enabled us to assess the consistency of our findings across simulated datasets and mitigate the impact of outliers or sampling bias.

### Supplemental Summary Statistics

**Table 1.**
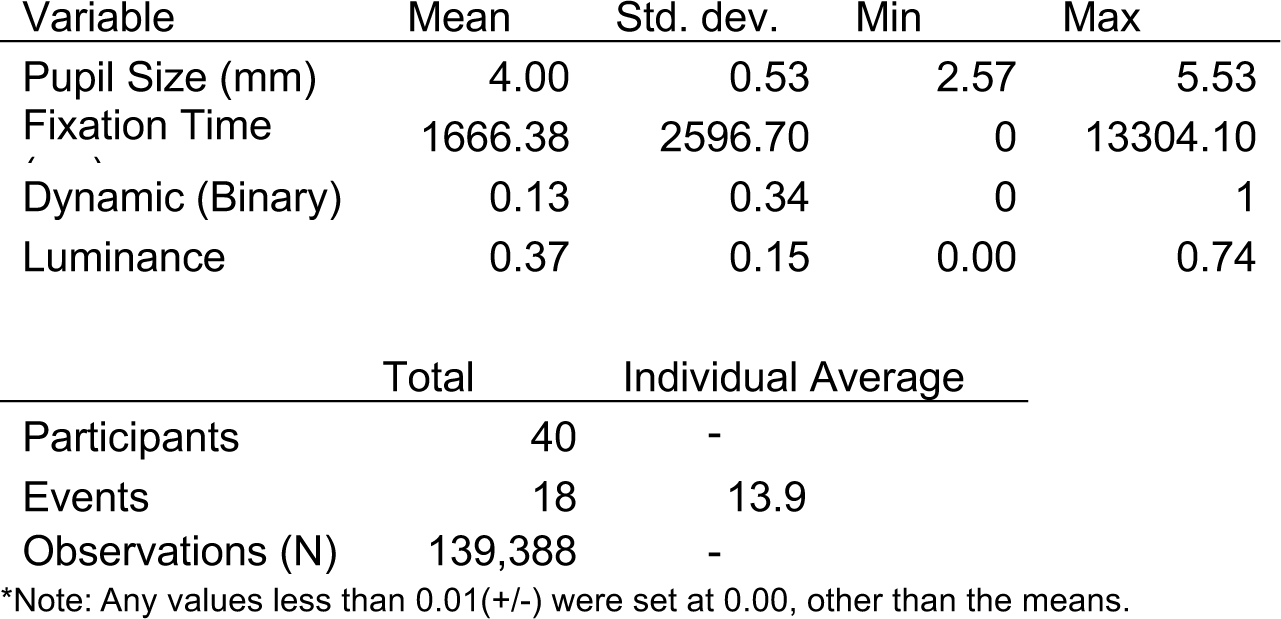
Summary of Variables Used in the First Stage Model.

Summary of Variables Used in the First Stage Model
| Variable | Mean | Std. dev. | Min | Max |
| --- | --- | --- | --- | --- |
| Pupil Size (mm) | 4.00 | 0.53 | 2.57 | 5.53 |
| Fixation Time | 1666.38 | 2596.70 | 0 | 13304.10 |
| Dynamic (Binary) | 0.13 | 0.34 | 0 | 1 |
| Luminance | 0.37 | 0.15 | 0.00 | 0.74 |

|  | Total | Individual Average |
| --- | --- | --- |
| Participants | 40 | - |
| Events | 18 | 13.9 |
| Observations (N) | 139,388 | - |

\*Note: Any values less than 0.01(+/-) were set at 0.00, other than the means.

**Table 2^12,15^.** Summary of Variables Used in the Second Stage Model.

Summary of Variables Used in the Second Stage Model
| Variable | Mean | Std. dev. | Min | Max |
| --- | --- | --- | --- | --- |
| Pupil Size (mm) | 4.00 | 0.47 | 3.13 | 5.09 |
| VABS Composite Score | 105.25 | 11.27 | 83 | 140 |
| Social PRI (ms) | -1.51x10<sup>-5</sup> | 0.00 | 0.00 | 0.00 |
| Non-Social PRI | 1.16x10<sup>-5</sup> | 0.00 | 0.00 | 0.00 |
| Average Fixation Time (ms) | 1525.34 | 795.85 | 148.69 | 3972.68 |
| Age (months) | 12.05 | 4.72 | 6 | 22 |

|  | Total |
| --- | --- |
| Observations (N) | 40 |

\*Note: Any values less than 0.01(+/-) were set at 0.00, other than the means.

**Table 3.**
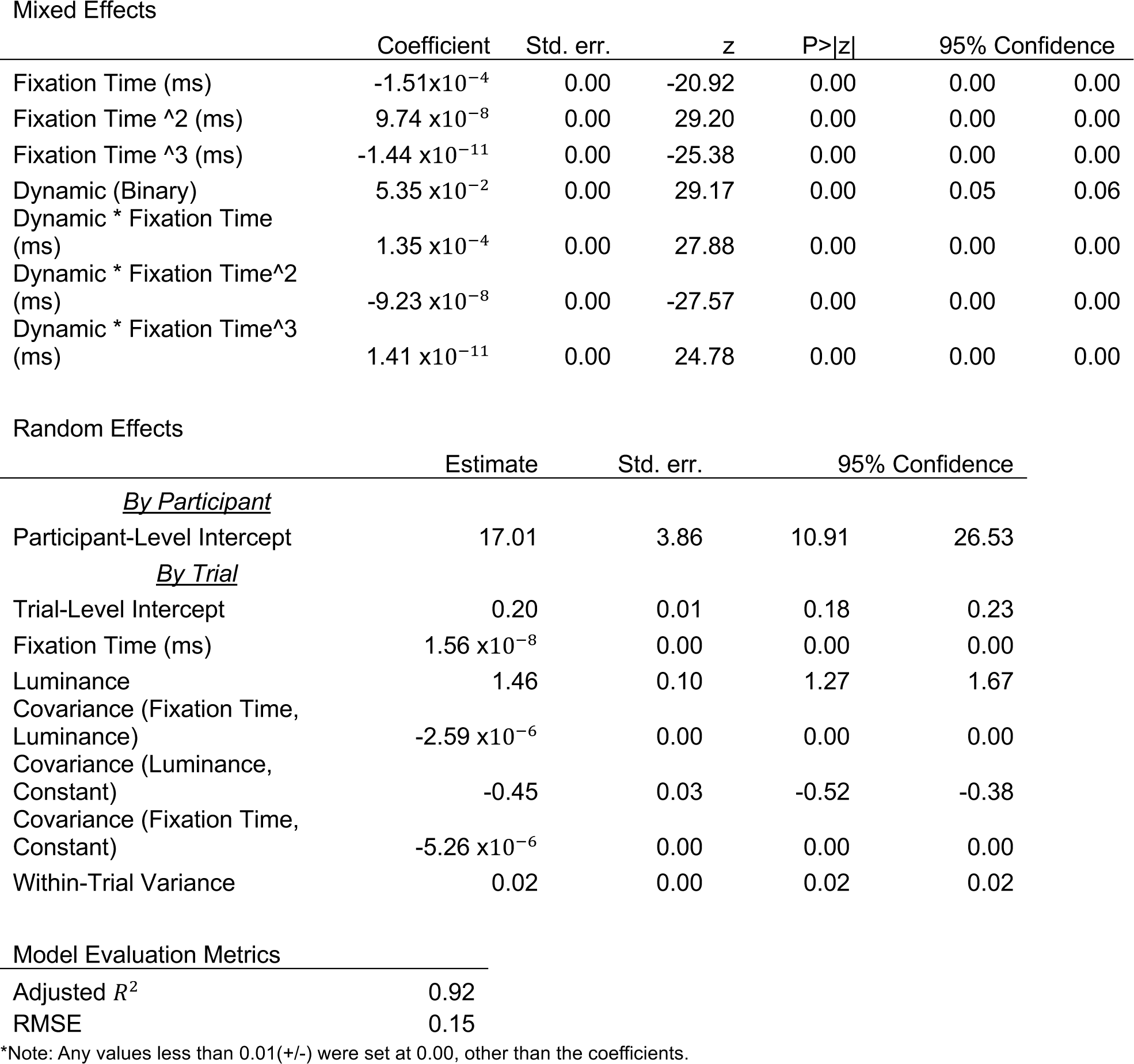
First Stage Regression Results: Multi-Level Mixed Effects Model.

Mixed Effects
|  | Coefficient | Std. err. | z | P>\|z\| | 95% Confidence |  |
| --- | --- | --- | --- | --- | --- | --- |
| Fixation Time (ms) | -1.51x10<sup>-4</sup> | 0.00 | -20.92 | 0.00 | 0.00 | 0.00 |
| Fixation Time ^2 (ms) | 9.74 x10<sup>-8</sup> | 0.00 | 29.20 | 0.00 | 0.00 | 0.00 |
| Fixation Time ^3 (ms) | -1.44 x10<sup>-11</sup> | 0.00 | -25.38 | 0.00 | 0.00 | 0.00 |
| Dynamic (Binary) | 5.35 x10<sup>-2</sup> | 0.00 | 29.17 | 0.00 | 0.05 | 0.06 |
| Dynamic * Fixation Time (ms) | 1.35 x10<sup>-4</sup> | 0.00 | 27.88 | 0.00 | 0.00 | 0.00 |
| Dynamic * Fixation Time^2 (ms) | -9.23 x10<sup>-8</sup> | 0.00 | -27.57 | 0.00 | 0.00 | 0.00 |
| Dynamic * Fixation Time^3 (ms) | 1.41 x10<sup>-11</sup> | 0.00 | 24.78 | 0.00 | 0.00 | 0.00 |

Random Effects
|  | Estimate | Std. err. | 95% Confidence |  |
| --- | --- | --- | --- | --- |
| *<u>By Participant</u>* |  |  |  |  |
| Participant-Level Intercept | 17.01 | 3.86 | 10.91 | 26.53 |
| *<u>By Trial</u>* |  |  |  |  |
| Trial-Level Intercept | 0.20 | 0.01 | 0.18 | 0.23 |
| Fixation Time (ms) | 1.56 x10<sup>-8</sup> | 0.00 | 0.00 | 0.00 |
| Luminance | 1.46 | 0.10 | 1.27 | 1.67 |
| Covariance (Fixation Time, Luminance) | -2.59 x10<sup>-6</sup> | 0.00 | 0.00 | 0.00 |
| Covariance (Luminance, Constant) | -0.45 | 0.03 | -0.52 | -0.38 |
| Covariance (Fixation Time, Constant) | -5.26 x10<sup>-6</sup> | 0.00 | 0.00 | 0.00 |
| Within-Trial Variance | 0.02 | 0.00 | 0.02 | 0.02 |

Model Evaluation Metrics
|  |  |
| --- | --- |
| Adjusted $R^2$ | 0.92 |
| RMSE | 0.15 |

\*Note: Any values less than 0.01(+/-) were set at 0.00, other than the coefficients.

**Table 4.**
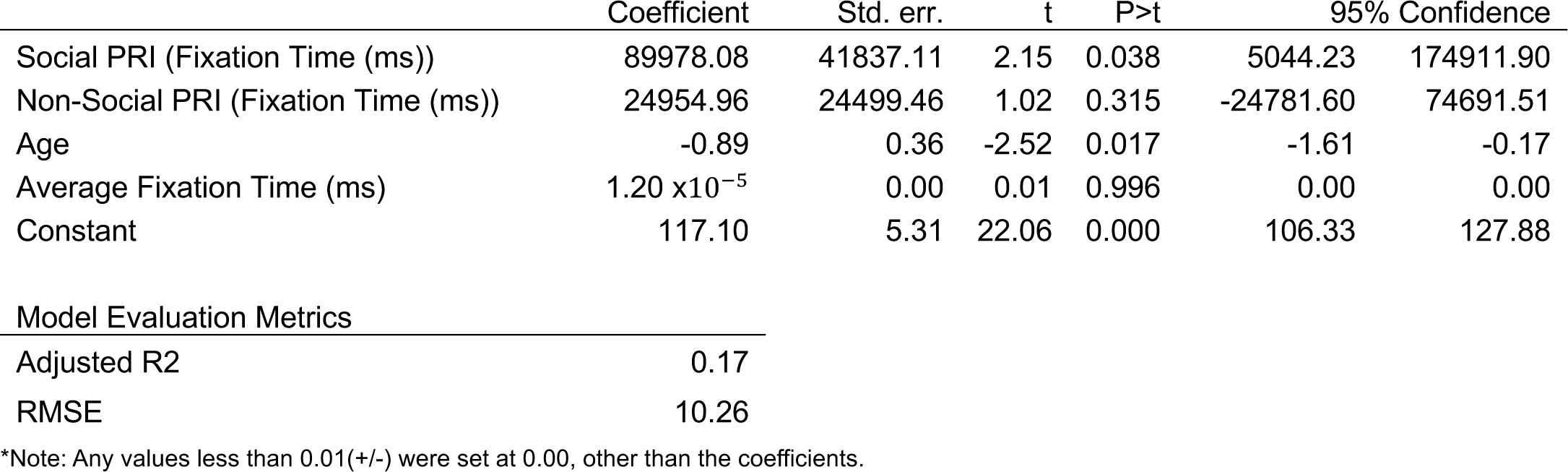
Second Stage Regression Results: Predict VABS Composite Scores per Participant.

|  | Coefficient | Std. err. | t | P>t | 95% Confidence |  |
| --- | --- | --- | --- | --- | --- | --- |
| Social PRI (Fixation Time (ms)) | 89978.08 | 41837.11 | 2.15 | 0.038 | 5044.23 | 174911.90 |
| Non-Social PRI (Fixation Time (ms)) | 24954.96 | 24499.46 | 1.02 | 0.315 | -24781.60 | 74691.51 |
| Age | -0.89 | 0.36 | -2.52 | 0.017 | -1.61 | -0.17 |
| Average Fixation Time (ms) | $1.20 \times 10^{-5}$ | 0.00 | 0.01 | 0.996 | 0.00 | 0.00 |
| Constant | 117.10 | 5.31 | 22.06 | 0.000 | 106.33 | 127.88 |

**Model Evaluation Metrics**
|  |  |
| --- | --- |
| Adjusted R2 | 0.17 |
| RMSE | 10.26 |

\*Note: Any values less than 0.01(+/-) were set at 0.00, other than the coefficients.

